# Nitric oxide-induced tyrosine nitration of TrkB impairs BDNF signaling and restrains neuronal plasticity

**DOI:** 10.1101/2022.05.07.491033

**Authors:** Caroline Biojone, Plinio C Casarotto, Cecilia Cannarozzo, Senem Merve Fred, Rosa Herrera-Rodríguez, Angelina Lesnikova, Mikko Voipio, Eero Castrén

**Affiliations:** Neuroscience Center, HiLife, University of Helsinki, Helsinki, Finland, 00790; University of Barcelona

**Keywords:** nitric oxide, TRKB, BDNF, plasticity, nNOS, nitration, visual cortex, ocular dominance

## Abstract

Nitric oxide has been long recognized as an important modulator of neural plasticity, but characterization of the molecular mechanisms involved - specially the guanylyl cyclase-independent ones - has been challenging. There is evidence that NO could modify BDNF-TRKB signaling, a key mediator of neuronal plasticity. However, the mechanism underlying the interplay of NO and TRKB remains unclear. Here we show that nitric oxide induces nitration of the tyrosine 816 in the TRKB receptor *in vivo* and *in vitro,* and that post-translational modification inhibits TRKB phosphorylation and binding of phospholipase Cγ1 (PLCγ1) to this same tyrosine residue. Additionally, nitration triggers clathrin-dependent endocytosis of TRKB through the adaptor protein AP2M and ubiquitination, thereby increasing translocation of TRKB away from the neuronal surface and directing it towards lysosomal degradation. Accordingly, inhibition of nitric oxide increases TRKB phosphorylation and TRKB-dependent neurite branching in neuronal cultures. *In vivo,* chronic inhibition of neuronal nitric oxide synthase (nNOS) dramatically reduced TRKB nitration and facilitated TRKB signaling in the primary visual cortex, and promoted a shift in ocular dominance upon monocular deprivation in the visual cortex - an indicator of increased plasticity. Altogether, our data describe and characterize a new molecular brake on plasticity, namely nitration of TRKB receptors.

**Significance statement:** We described the nitration of TRKB receptors at the tyrosine residue 816 as a new post-translational modification (PTM) that restrains the signaling of the neurotrophic factor BDNF in neurons. This new PTM leads to endocytosis and degradation of the TRKB receptors. Intriguingly, this mechanism is tonically active under physiological conditions *in vivo,* and it is important for restricting ocular dominance plasticity in the visual cortex. This mechanism directly links two major systems involved in brain plasticity, BDNF/TRKB and nitric oxide. Our data provides a model for *how* NO production from nNOS can compromise TRKB function, and for the effects of nNOS inhibitors promoting plasticity.

## Introduction

Nitric oxide (NO) is an important neuromodulator and, in the brain, it is mainly produced by the neuronal nitric oxide synthase (nNOS) (Huang et al., 1993), an enzyme that is tightly controlled by intracellular calcium levels and neuronal activity (Garthwaite et al., 1988; Steinert et al., 2011). NO has been shown to modulate several neuronal processes, such as neuronal intrinsic excitability (Steinert et al., 2011; Artinian et al., 2012), mobility of synaptic vesicles (Chenouard et al., 2020), GABA and glutamate release (Bradley and Steinert, 2016), expression of synaptic proteins in both pre- and postsynaptic neurons (Wang et al., 2005), and both long-term potentiation (LTP) or depression (LTD) (Schuman, 1995). In general, effects of NO are complex and bidirectional, depending on the brain area, concentration, and cellular environment, variables that directly determine the mechanism of action and signaling pathways recruited (Bradley and Steinert, 2016).

Being a gas, NO is a very peculiar messenger molecule: it is short-lived (0.5-5 seconds) and membrane permeable, and the local range of its effects can be estimated but not precisely determined, since the local environmental conditions (pH, presence of endogenous oxidants/antioxidants and scavengers, etc) can restrict its effects (Steinert et al., 2010; Radi, 2013). Those characteristics make the identification of NO’s signaling targets and understanding of its molecular effects a challenging task. Although many important NO-target proteins have been well characterized (Hardingham et al., 2013; Okamoto and Lipton, 2015), such as soluble guanylate cyclase (sGC) (Krumenacker et al., 2004), it is now clear that an unknown number of proteins that remain to be characterized are likely to be functionally modified by NO (Biojone et al., 2015). However, the identity and functional role of these target proteins has not been characterized.

TRKB is activated by binding of its ligand brain-derived neurotrophic factor (BDNF), which triggers dimerization and autophosphorylation of TRKB tyrosine residues and initiates signaling via PI3K-Akt, MAPK/ERK, and PLCγ pathways (Minichiello, 2009). Of interest, the tyrosine residue at position 816 (Y816), that serves as a PLCγ docking site when phosphorylated, seems to be a potential target of nitration (Biojone et al., 2015). Thus, we aim at verifying if Y816 is nitrated in biological relevant conditions and if that post-translational modification affects TRKB signaling capability.

TRKB receptors are well-known orchestrators of plasticity in different levels (Umemori et al., 2018), and TRKB-mediated plasticity has been particularly well characterized in the visual cortex of mice, which prompted us to investigate the NO effects in that brain area. Of interest, experiments using the ocular dominance model (Cang et al., 2005) have revealed that intact TRKB signaling is needed for visual cortex plasticity induced by different experimental interventions (Lesnikova et al., 2020). Additionally, optogenetic activation of TRKB in parvalbumin-containing interneurons *per se* in the visual cortex of adult mice renders the circuitry more plastic, allowing the ocular dominance to shift according to the changes in the visual stimuli (Winkel et al., 2021). TRKB-induced juvenile like-plasticity in the visual cortex involves modulation of cortical excitatory/inhibitory balance (Winkel et al., 2021), a mechanism that has been long recognized as critical for the closure of critical periods of plasticity (Fagiolini and Hensch, 2004; Hensch, 2005). Although endogenous NO signaling has also been shown to modulate E/I balance in the visual cortex in 18-25 days young rats (Le Roux et al., 2009) (age compatible with the beginning of the critical period (Fagiolini et al., 1994)), and mediate lateral inhibition to neighboring columns in the immature somatosensory cortex (Shlosberg et al., 2012), the potential effects of NO on plasticity of the adult visual cortex remains to be investigated.

In this study, we show *in vitro* and *in vivo* that under physiological conditions, tonic production of NO by nNOS restrains TRKB signaling through nitration at the Y816. Nitration triggers TRKB removal from the neuronal surface through coupling with clathrin endocytic adaptor AP-2 and drives TRKB to lysosomal degradation through ubiquitination. TRKB signaling in the visual cortex can be derepressed by systemic inhibition of nNOS, which then allows the reopening of the critical period of plasticity in that brain area. Altogether, our data suggests that NO works as a molecular brake of TRKB-induced plasticity in the visual cortex.

## Methods

### Drugs and reagents

NPA (n^ω^-propyl-L-arginine hydrochloride, #1200, Tocris), ODQ (1*H*-[1,2,4]Oxadiazolo[4,3-*a*]quinoxalin-1-one, #0880, Tocris), SNP (sodium nitroprusside dihydrate, #71778, Sigma-Aldrich), L-arginine (#A5006, Sigma-Aldrich), and recombinant human BDNF (#450-02, PeproTech) were used. SNP was freshly prepared immediately before each experiment and protected from light. BDNF was diluted in sterile PBS, ODQ was diluted in DMSO:miliQ water (1:1000), and the other drugs were diluted in sterile miliQ water. In cell culture experiments, drugs were added directly to the media, and the vehicle did not exceed 0.01% of media volume.

### Neuronal culture preparation, stimulation, and sample collection

Neurons from hippocampal or cortical brain tissue were collected from Wistar rats at embryonic day 17-18, as previously described (Sahu et al., 2019), and seeded in 1.9 cm^2^ wells (125,000/well), in poly-L-lysine coated plates (for protein analysis) or glass coverslips (80,000/well for immunohistochemistry experiments). Exceptionally for TRKB on cell surface experiment, neurons were seeded in 96-HB TC ViewPlate (Perkin Elmer), at 60,000/well. Cultures were maintained in Neurobasal media supplemented with 1% B-27, 1% penicillin/streptomycin, and 1% L-glutamine at 37°C, in 5% CO_2_ humidified incubator for 7 days before drug stimulation started. Drug treatment was added at diverse concentration and duration (which are informed in the legend of respective figures), then the media was removed and neurons were washed with PBS once, then the samples were collected for immunohistochemistry and surface experiments by fixing with 4% PFA 20 min RT, or for protein analysis (ELISA and WB experiments) by adding NP lysis buffer (20 mM Tris-HCl, 150 mM NaCl, 50 mM NaF, 1% Nonidet-40, 10% glycerol) supplemented with 2 mM Na3VO4 and cOmplete inhibitor mix (Roche) followed by harsh agitation at 4°C 30 min and centrifugation (4°C, 15,000 g, 10 min).

### Analysis of TRKB phosphorylation and nitration in brain tissue

18 weeks old male C57BL/6J RccHsd mice acquired from Envigo (Harlan Labs, UK) were used in these experiments. Mice were group housed under standard conditions (temperature 22°C, 12h light cycle) with food and water *ad libitum.* NPA 1 mg/kg was delivered by peroral in the drinking water for 3 weeks. The volume of water ingested was monitored and used to adjust the drug concentration in the water and consequent dosage. After the treatment, the mice were euthanized with CO_2_ inhalation and decapitation, and the visual cortex was collected, frozen in dry ice, and stored at −70°C. Samples were homogenized in NP lysis buffer (20 mM Tris-HCl, 150 mM NaCl, 50 mM NaF, 1% Nonidet-40, 10% glycerol) supplemented with 2 mM Na_3_VO_4_ and cOmplete inhibitor mix (Roche) and centrifugation (4°C, 15,000 g, 10 min). Then, the pTRKB, nitroTRKB, and total TRKB were assessed by sandwich ELISA (described below). All experiments were conducted following International guidelines for animal experimentation and were approved by the County Administrative Board of Southern Finland (License number: ESAVI/7551/04.10.07/2013). All efforts were made to minimize animal suffering.

### Nitration in N2a cell line

N2a cells were maintained in DMEM plus 10% FBS, and transfected with Lipofectamine 2000 (Thermo Fisher), according to manufacturer’s instructions, to express either WT or mutated GFP-tagged TRKB (mutation substitutes tyrosine, at position 816, for phenylalanine). Briefly, lipofectamine was incubated with the plasmid (2.5 μl:500 ng, in Opti-MEM) for 15 min RT, and added directly to the cell media at final concentration of 0.5% (v/v). 24h later, the medium was replaced by serum-free DMEM and the cells were stimulated with L-arginine 1.5 mM for 15 min. Protein samples were collected in NP lysis buffer (similarly as described above for neuronal cultures), and TRKB nitration was assessed by sandwich ELISA (described below).

### Immunoprecipitation and western blotting for detection of tyrosine nitration in TRKB

The samples were collected as above mentioned and acidified to dissociate protein complexes using HCl, and pH was restored using NaOH. Then, immunoprecipitation was done by incubation with mouse anti-nitroY antibody (#sc32757, Santa Cruz) overnight 4°C, followed by Sepharose G incubation (2h RT under agitation). After centrifugation (2 min, 2000g), the pellet was washed with NP lysis buffer twice and prepared for SDS-PAGE by heating in 2X Laemmli buffer for 5 min at 95° C. Electrophoresis was carried out using NuPAGE 4-12% Bis-Tris Protein polyacrylamide gels (#NP0323BOX, Invitrogen). After the electrophoresis, the samples were transferred to PVDF membrane and incubated in rabbit anti-TRK antibody (sc-11, Santa Cruz Biotechnology) diluted 1:1000 in 3% BSA/TBST overnight at 4° C. The membrane was subsequently washed and incubated in anti-rabbit HRP-conjugated antibody (1:10,000, BioRad) for 1 hour RT. The bands were visualized using Pierce™ ECL Plus western blotting substrate (#32132, Thermo Fisher Scientific).

### ELISA for detection of PTM in TRKB and analysis of protein interaction

ELISA was used throughout different experiments, the antibodies and dilutions were substituted accordingly depending on the aim of the assay, but the same standard protocol and buffers were maintained. For analysis of endogenous TRKB interaction with AP2M, PLCγ1, and ubiquitin, goat anti-TRKB 1:500 (#AF1494, R&D systems) was used as primary antibody; anti-AP2M (1:2000, #sc-515920, Santa Cruz) or anti-PLCγ1 (1:2000; #5690, Cell Signaling) or anti-ubiquitin (1:1000, #sc-8017, Santa Cruz) were used as secondary antibodies. For analysis of Y816 TRKB phosphorylation, goat anti-TRKB 1:500 (#AF1494, R&D systems) was used as primary, and rabbit anti-phospho TRK 1:1000 (#4168, Cell Signaling) as a secondary antibody. For analysis of TRKB nitration, mouse anti-nitroY 1:500 (#sc32757, Santa Cruz) and goat anti-TRKB 1:1000 (#AF1494, R&D systems) were used as primary and secondary, respectively. For analysis of total TRKB, goat Ab against the extracellular domain of TRKB 1:500 (#AF1494, R&D systems) was used as primary, and rabbit Ab against the intracellular domain of TRKB 1:2000 (#92991, Cell Signaling) as a secondary antibody. Briefly, flat-bottom white plates (OptiPlate 96F-HB, Perkin Elmer) were coated overnight with primary antibody diluted in carbonate buffer pH 9.7 (25 mM sodium bicarbonate, 25 Mm sodium carbonate) at 4°C under agitation. Following blockade with 5% BSA in PBST for 2h at RT, the samples (lysate from neuronal culture) were incubated overnight under agitation at 4°C. The plates were washed (3x with PBST) followed by incubation with the secondary antibody overnight at 4°C under agitation. Following a washing step (3x PBST), the plates were incubated with 1:5000 HRP-conjugated tertiary antibody against the host of the secondary antibody for 2h at RT. Plates were washed, and the luminescence from the HRP activity was detected by incubation with ECL (Thermo Fisher Scientific) on a plate reader (Varioskan Flash, Thermo Scientific). Unspecific signal (average luminescence from 8-10 wells in which the samples were omitted) was assessed in every assay. Data was processed by subtracting nonspecific signal from sample signal, and then expressed as percentage of the control (vehicle treated group).

### C-terminal TRKB peptide design and analysis of its interaction with AP2M and PLCγ1

The synthetic peptides were designed based on the last 26 amino acid residues from C-terminal full length TRKB from rat (UniProt ID: Q63604) and ordered from Genscript, USA. Additionally, a biotin tag was added at the amino terminal portion to facilitate detection. Peptides were synthesized to contain a phosphorylated or nitrated tyrosine residue at position equivalent to Y816 (shown in bold) in the TRKB full length sequence. The amino acids sequences for the peptides are as follow:

control peptide: biotin-RKNIKNIHTLLQNLAKASPV**<u>Y</u>**LDILG;
phosphorylated peptide: biotin-RKNIKNIHTLLQNLAKASPV**[<u>pY</u>]**
LDILG; nitrated peptide: biotin-RKNIKNIHTLLQNLAKASPV**[<u>YNO2</u>]**LDILG.

Analysis of interaction between TRKB C-term peptides and AP2M or PLCγ1 was done by ELISA following the protocol above mentioned with minor modifications. Briefly, AP2M or PLCγ1 were pulled down from untreated neuronal cultures by coating OptiPlate overnight with antibodies (either anti-AP2M, #sc-515920, Santa Cruz, or anti-PLCγ1, #5690, Cell Signaling) diluted 1:500 in carbonate buffer, pH 9.8. Then, the plate was blocked in 5% BSA/PBST for 2h RT, and incubated with lysate from neuronal culture overnight at 4°C. C-terminal TRKB peptides (ctrl, phospho, or nitropeptide) diluted in 5%BSA/PBS were added at 1 μg per well and incubated overnight at 4°C. After washing with PBST, streptavidin-conjugated HRP (1:10,000 in 5% BSA/PBST) was added for 1h at RT, and HRP activity was measured by Varioskan plate reader, in the presence of ECL (Thermo Fisher Scientific).

### Model generation and solvent accessibility of nitrated TRKB C-terminal

The model of mouse TRKB.C-terminal (the last 26aa residues in C-terminal) was generated in the RaptorX server (Källberg et al., 2012), using wild-type (WT) sequence. The YNO2 was inserted to Y816 C3 (YNO2) using PyTMs plug-in for PyMOL (Warnecke et al., 2014), and the models were aligned WT and YNO2 for comparison, and the root-mean-squared deviation (RMSD) between the backbone carbon structure calculated by the server. The relative solvent accessibility per residues was calculated using PyMOL (v2.0 Schrödinger, LLC), and normalized by WT values.

### Immunostaining for colocalization of Rab7 or Lamp1 with TRKB

Hippocampal neurons were cultured in plates with poly-L-lysine coated glass coverslips (80000 cells/13 mm diameter coverslips). After vehicle (MilliQ) and SNP treatment for 30 min and 60 min, 7 DIV neurons were fixed in 4% paraformaldehyde in PBS for 20 min at RT. Following several washes in PBS, coverslips were incubated in the blocking buffer (5% donkey serum, 1% BSA, 0.1% gelatin, 0.1% Triton X-100, 0.05% Tween-20 in PBS) for 1 h at RT. Primary antibodies, rabbit anti-RAB7 (1:500, #D95F2, Cell Signaling) or mouse anti-Lamp1 (1:200, #sc-20011, Santa Cruz), and goat anti-TRKB (1:1000, #AF1494, R&D Systems) were diluted in the primary antibody buffer (1% BSA, 0.1% gelatin in PBS), and the coverslips were incubated ON at 4 °C under agitation. After brief washes in PBS, coverslips were incubated in Alexa Fluor–conjugated secondary antibodies (1:1000, Thermo Fisher Scientific) for 45 min at RT. Donkey anti-goat-647 was used for labeling TRKB and donkey anti-rabbit or mouse Alexa-568 for RAB7 and Lamp1, respectively. Final washes in PBS were followed by a brief wash in MilliQ and the coverslips were mounted in Dako Fluorescence Mounting Medium (S3023, Dako North America, Inc.). Imaging was performed at 647 and 568 channels with a Zeiss LSM710 confocal microscope, 63x oil objective at 1024×1024 pixel resolution. At least 8-20 Z-stack steps were acquired with 0.40 μm intervals. Mander’s coefficient was calculated in ZEN Imaging software (Zeiss, Germany) and used to assess the colocalization of Rab7:TRKB or Lamp1:TRKB in the soma.

### Sholl analysis of neurite branching

Cortical neurons were incubated with ANA12 0.1 μM or veh for 15 min, then with veh or NPA 0.4 ηM for 30 min, once a day for 3 days, at 8, 9, 10 DIV. 24h after the last treatment (11 DIV), the neurons were incubated with MgCl2 10 μM for 1h to prevent excitotoxicity and then transfected to express mCherry, using lipofectamine according to manufacturer’s instructions (similarly to the protocol described above in N2a experiment). The culture medium was replaced, 90 min after transfection, by fresh Neurobasal medium supplemented only with L-glu, B27 and penicillin/streptomycin. Next day, the coverslips were fixed with PFA 4% for 20 min, blocked, and counterstained with Hoescht 1:10,000 for 10 min. Coverslips were washed in PBST (3x) and miliQ water once, then mounted in DAKO fluorescence mounting media (S3023, Dako North America, Inc.). Imaging was acquired in Zeiss LSM700 confocal microscope, 25x oil objective at 1024×1024 pixel resolution, using 405 ηm (for Hoescht) and 568 ηm (mCherry) channels. At least 10 Z-stack steps were acquired. Confocal pictures were analysed in ImageJ (Fiji) software, by compiling the 568ηm z-stacks, setting the threshold automatically (Shanbhag threshold, available in the software), setting the center of the soma manually, and then counting the branching intersections automatically with built-in Sholl analysis tool (Ferreira et al., 2014). Sholl circle interval was set to 5 μm and the intersections were counted up to 100 μm of distance from the soma.

### Quantitation of TRKB on neuronal surface

The samples were collected as above mentioned and the levels of TRKB in the cell surface were determined as described in the literature (Zheng et al., 2008, Fred et al 2019). Briefly, the fixed neurons were blocked (5% BSA in PBS, 1h RT), the wells were incubated with anti-TRKB (R&D Systems, #AF1494, 1:1000 in blocking buffer) overnight at 4°C under agitation. After washing with PBS, the samples were incubated with HRP-conjugated anti-goat (1:5000 in blocking buffer) for 1h at RT. Finally, the cells were washed with PBS (4x for 10 min), and the chemiluminescent signal generated by reaction with ECL (Thermo Fisher Scientific) was analyzed in the Varioskan plate reader.

### Analysis of plasticity in the visual cortex induced by NPA

Plasticity in the visual cortex was investigated by depriving the left eye of visual stimulation for one week, and then later analyzing the compensatory adaptative response of the visual cortex to the visual input. For this experiment, 10 weeks old female C57BL/6J RccHsd mice were acquired from Envigo (Harlan Labs, UK) and group housed under standard conditions (temperature 22°C, 12h light cycle) with food and water *ad libitum.* Monocular deprivation started after spontaneous closure of critical period of plasticity in visual cortex (Lehmann and Löwel, 2008), *i.e.* when mice were 18 weeks old. NPA treatment 1 mg/kg (Montezuma et al., 2012) was delivered continuously in the drinking water for 4 weeks (bottles were changed twice a week), starting 3 weeks prior to the monocular deprivation. The volume of water ingested was monitored and used to adjust the drug concentration in the water and consequent dosage. The time course of the experimental procedures (skull preparation, repeated imaging of intrinsic signal, drug treatment and monocular deprivation) is shown in fig. 4A. All experiments were conducted following International guidelines for animal experimentation and were approved by the County Administrative Board of Southern Finland (License number: ESAVI/7551/04.10.07/2013). All efforts were made to minimize animal suffering.

**Fig 1:**
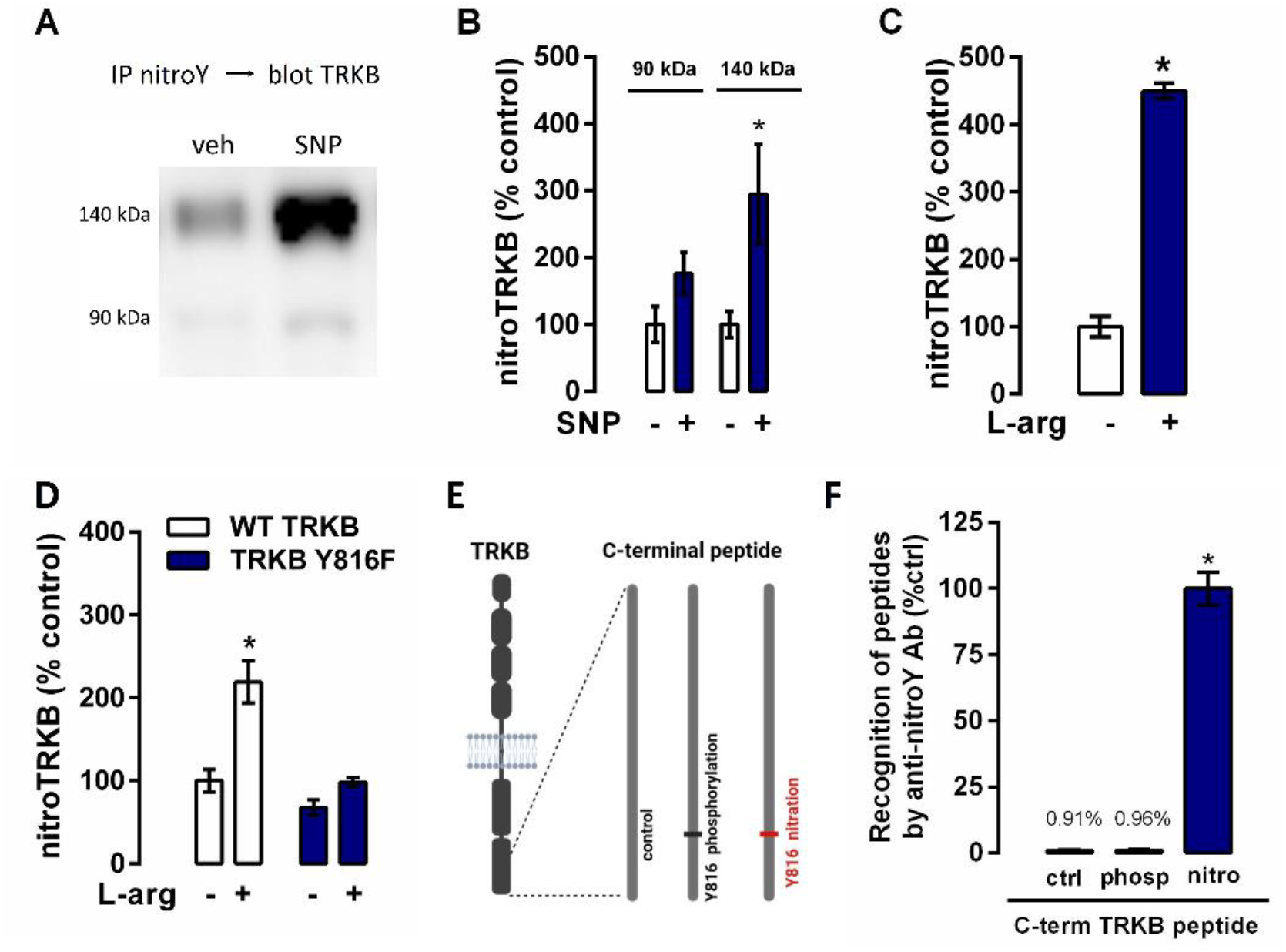
NO induces nitration of Y816 residue in TRKB receptors, *in vivo* and *in vitro.* (**A**) 8 DIV cortical neurons in culture were treated with NO donor sodium nitroprusside (SNP) 10 μM for 30 min, protein complexes were dissociated by acidification, pulled down with anti-nitroY antibody and detected by western blotting with anti-TRKB (representative samples from n=6/group. 140 and 90 kDa bands correspond to glycosylated and non-glycosylated forms of TRKB, respectively). (**B**) Quantitation of the experiment shown in A. (**C**) TRKB nitration was found also in 7 DIV hippocampal neurons treated with 1.5 mM L-arginine for 15 min, and analyzed by sandwich ELISA (n=6/group). (**D**) Y816F substitution renders TRKB insensitive to nitration induced by L-arginine (1.5 mM, 15 min), in N2a cells transfected to express WT (white columns) or Y816F mutated (blue columns) TRKB (n=8/group). (**E**) Synthetic peptides were designed based on the C-terminal sequence of TRKB, and nitration or phosphorylation were added to the intracellular domain at position Y816. Those peptides were used to validate antibody specificity (shown in F). (**F**) Anti-nitroY specifically pulled down nitrated TRKB C-terminal peptides but not phosphorylated or control peptides (n=5/group). Data was analyzed by unpaired T test (C), one-way ANOVA (F), or two-way ANOVA (B, D, E), and presented as mean±sem. *p<0.05.

**Fig 2:**
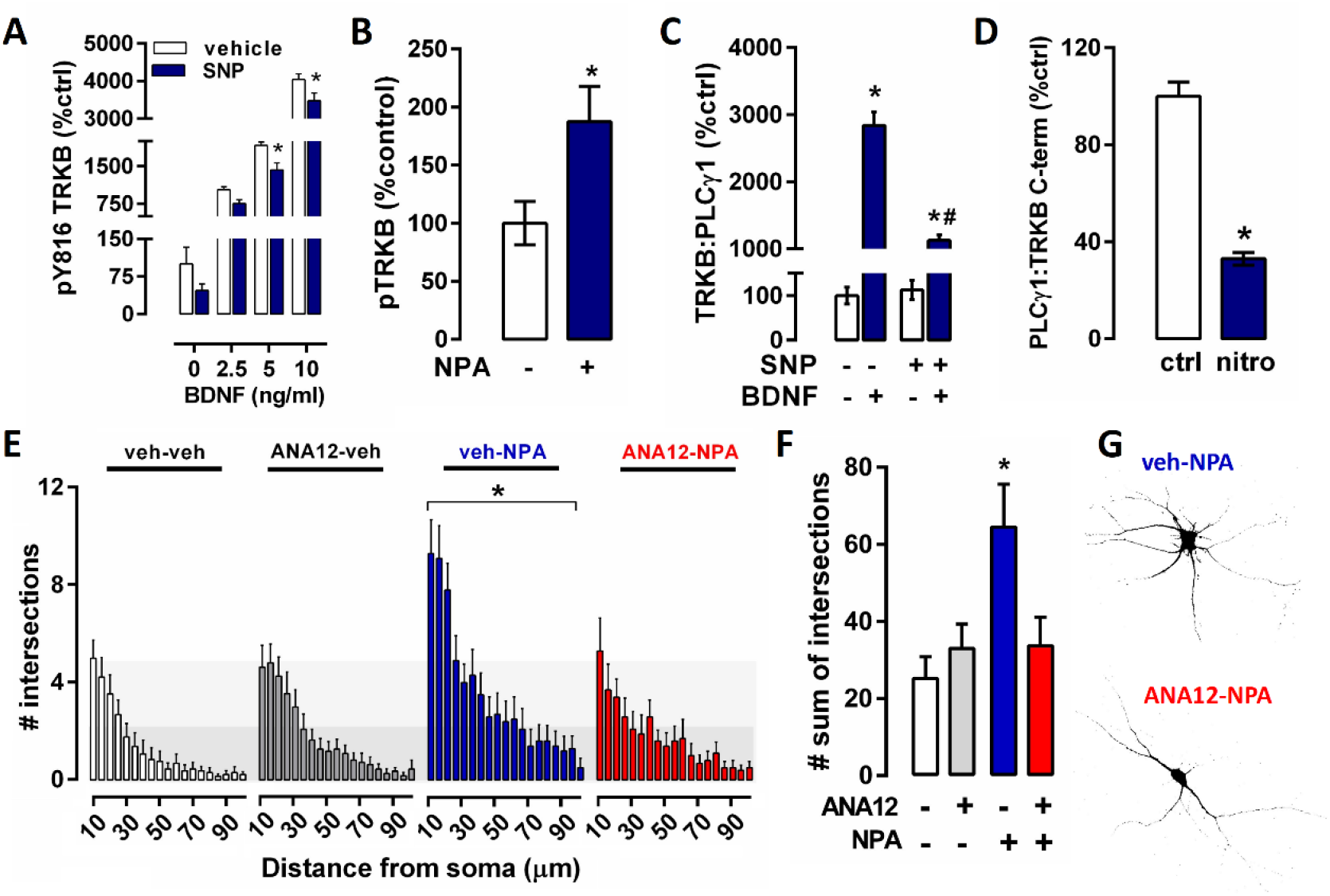
Nitration restricts TRKB phosphorylation and TRKB-PLCγl interaction. (**A**) In 7 DIV hippocampal neurons, BDNF-induced TRKB-phosphorylation is reduced when neurons are pretreated with NO donor sodium nitroprusside (SNP 10 μM, 15 min, n=5-6/group). (**B**) nNOS inhibitor n^ω^propyl-L-argininne (NPA 0.4 ηM, 30 min, n=8/group) *per se* increases TRKB phosphorylation. (**C**) SNP reduces BDNF-induced interaction between endogenous TRKB and PLCγ1 (co-IP, n=6/group) in hippocampal neurons. (**D**) There is reduced interaction between endogenous PLCγ1 and the synthetic 26 aa peptide corresponding to TRKB C-terminal tail nitrated at position equivalent to Y816, compared to the unmodified control peptide (n=6/group). (**E, F, G**) NPA 0.4 ηM (incubated for 30 min daily for 3 days, n=10-13/group) induced neuritogenesis in cortical neurons in culture, evaluated by Sholl analysis. Pretreatment with TRKB antagonist ANA12 (0.1 μM, 15 min) abolished the NPA effect. (**E**) Number of branching intersections at various distances from the cell body quantified every 5 μm up to 100 μm. (**F**) Total number of intersections per neuron, irrespective of the distance from the soma. (**G**) Representative pictures from samples analyzed in F and G. Data was analyzed by unpaired T test (B, D) or two-way ANOVA (A, C, E, F), and presented as mean±sem. *p<0.05.

**Fig 3:**
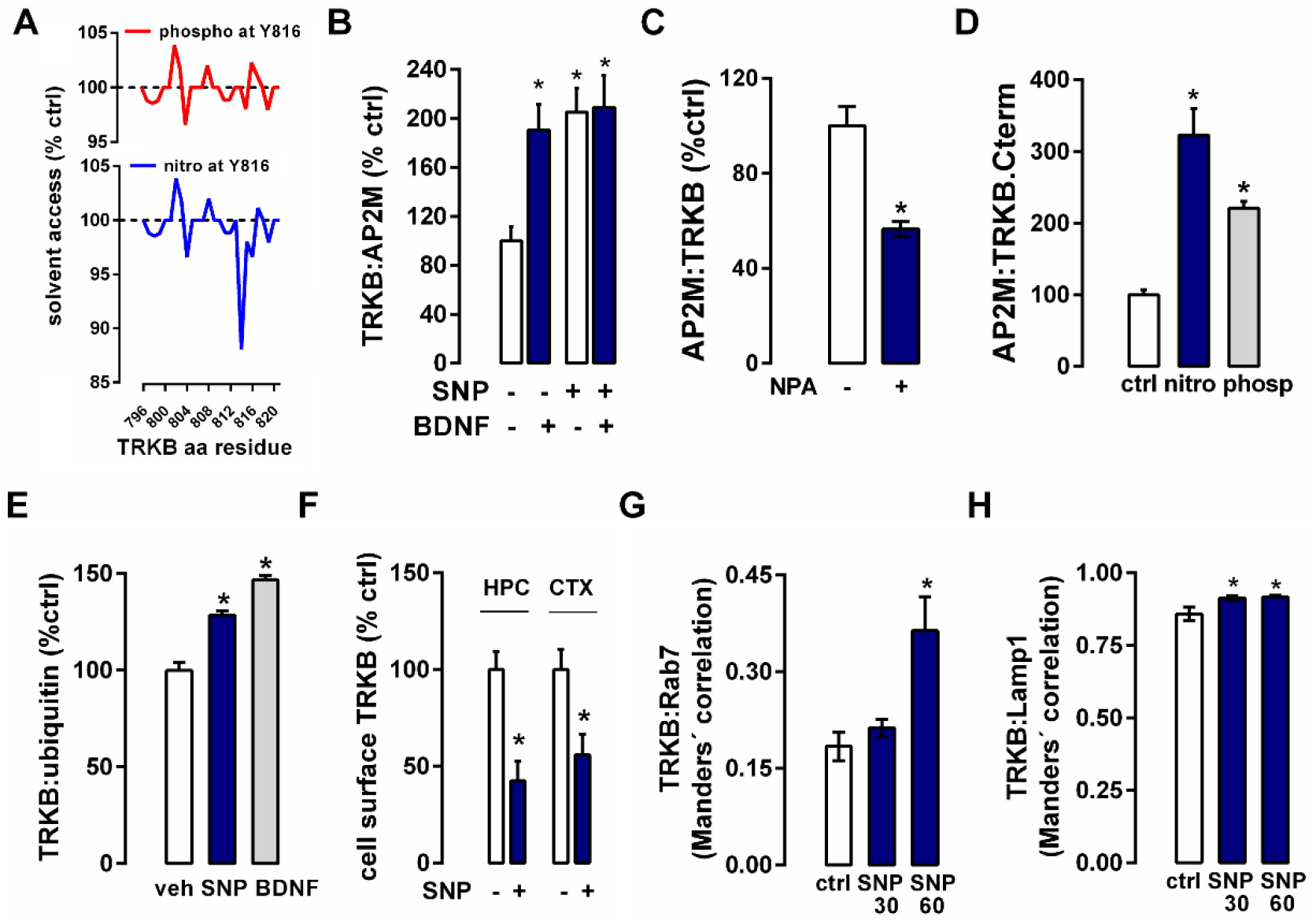
Nitration modifies TRKB conformation, and triggers its endocytosis and degradation. (**A**) Effect of Y816 nitration (bottom panel, blue line) or phosphorylation (upper panel, red line) upon solvent accessibility of TRKB C-terminal portion (shown in the x axis), normalized by control (non-nitrated and non-phosphorylated, represented by the dashed line at 100%), calculated using PyMOL software. (**B**) NO donor SNP (10 μM 15 min, n=5-6/group) increases while (**C**) nNOS inhibitor NPA (0.4 ηM, 30 min, n=6/group) decreases the interaction (co-IP) between endogenous TRKB and the adaptor protein complex AP2M in hippocampal neurons. (**D**) The interaction between endogenous AP2M and the synthetic peptide (depicted in fig 1E) either nitrated or phosphorylated at position equivalent to Y816, is increased compared to the control peptide (n=15-16/group). (**E**) SNP increases ubiquitination of TRKB within 15 min (n=8/group) and (**F**) decreases TRKB on the cell surface (30 min, n=8-11/group). (**G**)Co-localization of TRKB with the late endocytic marker Rab7 (n=9-13/group) and with(**H**)the lysosomal marker Lamp1 (n=15/group) was increased at 30 and 60 min after SNP treatment. Data was analyzed by unpaired T test (C), Kruskal-Wallis test (D, G), one-way ANOVA (E, H), two-way ANOVA (B, F), and presented as mean±sem. *p<0.05.

**Fig 4:**
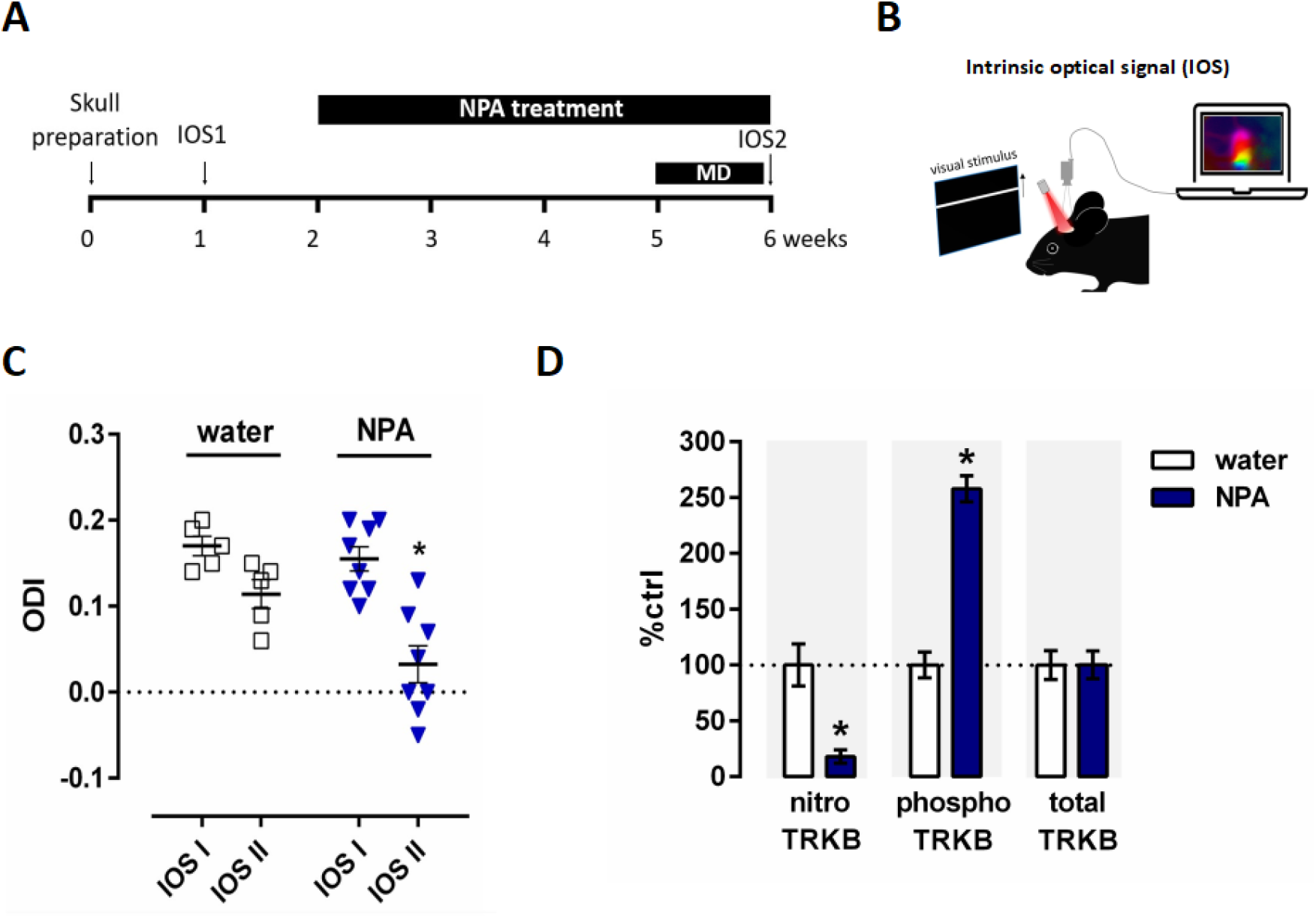
nNOS inhibitor NPA (1 mg/kg p.o.) facilitates plasticity in the adult mouse visual cortex. (**A**) Scheme showing the time course of the experimental procedures. MD: monocular deprivation. IOS: intrinsic optical signal. (**B**) Depiction of IOS acquisition: visual stimulus (wide horizontal bar moving upwards) was presented to either left or right eye, in a screen, and the IOS of the visual cortex was acquired under red light stimulation to the skull. (**C**) Combination of NPA treatment and MD triggers a shift in the ocular dominance towards the non-deprived eye (n=5-8/group). The Ocular Dominance Index (ODI) values range from −1 to +1: positive values correspond to a contralateral bias, negative ones indicate ipsilateral bias, while ODI values of 0 show that ipsilateral and contralateral eyes are equally strong. (**D**) NPA treatment (1 mg/kg p.o. during 3 weeks) reduced nitration and increased phosphorylation in TRKB in the visual cortex of naive mice. Expression of TRKB (total TRKB) was not changed by NPA treatment. Data was analyzed by two-way ANOVA, *p<0.05. Horizontal and vertical bars represent mean±sem respectively, and scattered points represent individual subjects.

#### Skull preparation

The Transparent Skull was prepared as previously described (Steinzeig et al., 2017). Briefly, animals were anesthetized with a mix of 0.05 mg/kg Fentanyl, 5 mg/kg Midazolam, and 0.5 mg/kg Medetomidine i.p., and placed on the stereotaxic frame with the body temperature maintained at 37° C. A mixture of Lidocaine and adrenaline (20 mg/ml, Orion Pharma, Finland) was locally injected subcutaneously on the head and the scalp was removed. The periosteum was gently scratched away from the skull and a cotton swab soaked in acetone was rapidly passed to clean the skull from fat. The surface of the skull was covered with a thin layer of cyanoacrylate glue (Loctite 401, Henkel, Germany) followed by two layers of acryl, to preserve the transparency. The acryl was prepared by stirring acrylic powder (EUBECOS, Germany) with methacrylate liquid (Densply, Germany) until a nail-polish consistency was reached. The transparent skull was left to dry overnight, after the mice were injected s.c. with 5 mg/kg Carprofen (ScanVet, Nord Ireland) for postoperative analgesia and injected i.p. with a wake-up mix composed by: 1.2 mg/Kg Naloxone (Orpha-Devel Handels und Vertriebs GmbH, Austria), opioid receptor antagonist; 0.5 mg/Kg Flumazenil (Hameln, Germany), GABA-A receptor antagonist; 2.5 mg/Kg Atipamezole (Vet Medic animal Health Oy, Finland), adrenergic receptor antagonist; diluted in saline. The next day, isoflurane at 4% was used to induce the anaesthesia and it was maintained at 2% for the procedure. The acryl layer was polished, a metal head holder was first glued on the skull, carefully keeping the area of interest at the center of the holder, and then fixed with a mixture of cyanoacrylate glue and dental cement (Densply, Germany). Finally, transparent nail polished (#72180, Electron Microscopy Sciences) was applied inside the metal holder.

#### Monocular deprivation

Isoflurane at 4% was used to induce the anaesthesia and then it was maintained at 2% until the end of the procedure. A drop of antibiotic eye gel (Isothal Vet 1%, Dechra, Canada) was applied on the left eye and the eye was closed with 3 mattress sutures. Antibiotic ointment (Oftan Dexa-Chlora, Anten, Finland) was applied on the sutured eye and Carprofen (5 mg/kg) was injected s.c. for postoperative analgesia. The monocular deprivation lasted 7 days, during which all animals were checked daily to prevent reopening of the eyes.

#### Optical imaging of the intrinsic signal

The changes in neurovascular coupling in the primary visual cortex of the right hemisphere were measured to assess visually-evoked cortical activity, as previously described (Steinzeig et al., 2017), by adapting a protocol developed by the laboratory of Dr. Stryker (Kalatsky and Stryker, 2003; Cang et al., 2005). The animals were anesthetized with 1.8% isoflurane with a 1:2 mixture of O2:air for 15 minutes and then maintained at 1.2% isoflurane for at least 10 minutes before starting the imaging session. Two sessions of imaging were performed: one before the beginning of the treatment administration (IOS I) and one after the 7th day of monocular deprivation (IOS II). Briefly, the animals were kept on a heating pad in front of and within 25 cm from the stimulus monitor. The head holder was fixed and the animal’s nose was aligned to the midline of the stimulus monitor. The visual stimulus was a 2° wide horizontal bar moving upwards with a temporal frequency of 0.125 Hz and a spatial frequency of 1/80 degree, displayed in the central part of a high refresh rate monitor (−15 to 5 degree azimuth, relative to the animal visual field) to stimulate the binocular part of the visual field. To acquire a map of the surface vascular pattern, the skull was illuminated with a green light (540±20 ηm). Then the camera was focused 600 μm below the pial surface and a red light (625±10 ηm) was used to record the intrinsic signal. The signal was recorded from one eye at the time, with the other eye covered by a patch. The continuous-periodic stimulation was synchronized with a continuous frame acquisition, frames were collected at a rate of 30 fps for 5 minutes and stored as a 512 x 512 pixel image. The intrinsic signal was extracted by using the analysis software package created in the laboratory of Stryker by Kalatsky, that performs Fourier decomposition on the cortical maps (Kalatsky and Stryker, 2003). The intrinsic response was measured as fractional changes in skull surface reflectance x 104 and its magnitude was used to calculate the activation of the visual cortex in the right hemisphere due to ipsilateral or contralateral eye stimulation. A low-pass filter (uniform kernel of 5 x 5 pixel) was applied to the ipsilateral magnitude map to smoothen it and the 30% of the peak response amplitude was set as threshold, to eliminate background noise and to define the area that produced the strongest response to the ipsilateral eye. The resulting map was used as a mask to select the binocularly responsive region of interest within the visual cortex. Cortical maps were calculated for both contralateral (C) and ipsilateral (I) eyes and the Ocular Dominance score was computed as (C–I)/(C+I). Finally, the Ocular Dominance Index (ODI) was calculated as the mean of the OD score for all responsive pixels (Cang et al., 2005). The ODI values range from −1 to +1: positive values correspond to a contralateral bias, negative ones indicate ipsilateral bias and ODI values of 0 show that ipsilateral and contralateral eyes are equally strong (Steinzeig et al., 2017).

### Statistical analysis

Data was analyzed preferentially by parametric tests (T test, one- and two-way ANOVA) to gain statistical power, unless the data presented lack of homoscedasticity or when variables were discrete, in which cases non-parametric tests were chosen (Kruskal-Wallis test). The statistical tests used in each particular experiment are described in the legend of figures, detailed statistical values (df, F, T, H, p) are described in the table 1. Differences were considered statistically significant when p<0.05. Statistical analysis and plots were made in GraphPad Prism 6 software. Data are presented as mean± SEM (column and error bars), and scattered symbols represent individual samples.

**Table 1:** Statistical analysis

| Figure | Test |
| --- | --- |
| 1B | Two-way ANOVA: Interaction: $F_{(1,20)}=1.856$, $p=0.1883$; MW: $F_{(1,20)}=1.856$, $p=0.1883$; SNP: $F_{(1,10)}=9.681$, $p=0.0055$. Bonferroni's multiple comparison test: $p=0.0098$ |
| 1C | Unpaired T test: $T_{10}=18.23$; $p<0.0001$ |
| 1D | Two-way ANOVA: Interaction: $F_{(1,28)}=8.31$, $p=0.0075$; L-arg: $F_{(1,28)}=23.68$, $p<0.0001$; mutation: $F_{(1,28)}=24.93$, $p<0.0001$. Bonferroni's multiple comparison test: $P<0.0001$ |
| 1F | One-way ANOVA: $F_{(2, 12)} = 250.1$, $p<0.0001$<br>Bonferroni's multiple comparison test: $P<0.0001$ |
| 2A | Two-way ANOVA: Interaction: $F_{(3,39)} = 2.182$, $p=0.10$; BDNF: $F_{(3,39)} = 398.4$; $p<0.0001$; SNP: $F_{(1,39)}= 19.73$, $p<0.0001$. Bonferroni's multiple comparison test: $P<0.012$ (vs ctrl). |
| 2B | Unpaired T test: $T_{14} = 2.458$; $p<0.0276$ |
| 2C | Two-way ANOVA: Interaction: $F_{(1,20)}= 61.88$; $p<0.0001$; SNP: $F_{(1,20)}= 60.11$; $p<0.0001$<br>BDNF $F_{(1,20)}= 293.8$; $p<0.0001$. Bonferroni's multiple comparison test: *$p<0.0001$ (vs veh-veh), #$p<0.0001$ (vs veh-BDNF). |
| 2D | Unpaired T test: $T_{10}=10.38$; $p<0.0001$ |
| 2E | Two-way ANOVA: Interaction: $F_{(54,720)}=1.897$, $p=0.0002$; distance: $F_{(18,720)}=48.46$, $p<0.0001$; treatments: $F_{(3,40)}=5.029$, $p=0.0047$. Tukey's multiple comparison test: $P<0.05$ (veh-NPA is different from every other group). |
| 2F | Two-way ANOVA: Interaction: $F_{(1,40)}=6.366$, $p=0.0157$; NPA: $F_{(1,40)}=6.817$, $p<0.0126$<br>ANA12: $F_{(1,40)}=2.259$, $p=0.1407$. Tukey's multiple comparison test: $P<0.05$ (veh-NPA is different from every other group). |
| 3B | Two-way ANOVA: Interaction: $F_{(1,19)}=4.482$, $p=0.0477$; SNP: $F_{(1,19)}=9.153$, $p<0.0070$; BDNF: $F_{(1,19)}=5.365$; $p=0.0319$. Bonferroni's multiple comparison test: $P<0.0384$ |
| 3C | Unpaired T test: $T_{10}=4.936$; $p=0.0006$ |
| 3D | Kruskal-Wallis: $H=32.33$; $p<0.0001$. Dunn's multiple comparison: $P<0.0001$ (vs ctrl) |
| 3E | One-way ANOVA: $F_{(2,21)}= 62.64$; $p<0.0001$. Bonferroni's multiple comparison test: $P<0.0001$ (vs ctrl) |
| 3F | Two-way ANOVA: Interaction: $F_{(1,33)}=0.4257$, $p=0.5186$; Brain area: $F_{(1,33)}=0.4296$, $p=0.5167$; Treatment: $F_{(1,33)}=24.21$, $p<0.0001$. Bonferroni's multiple comparisons test: veh x SNP: $p<0.02$ |
| 3G | Kruskal-Wallis: $H=6.383$; $p=0.0411$. Dunn's multiple comparison: $p=0.0317$ (vs ctrl) |
| 3H | One way-ANOVA: $F_{(2,42)}=4.927$; $p=0.0120$. Bonferroni's multiple comparison test: $P<0.0240$ (vs ctrl). |
| 4C | Two-way ANOVA: Interaction: $F_{(1,11)}=2.517$, $p=0.1409$; Time: $F_{(1,11)}=18.13$, $p=0.0013$; NPA: $F_{(1,11)}=10.02$, $p=0.0090$. Bonferroni's multiple comparison: $p=0.0013$ (NPA, IOS1 vs IOS2), $p=0.2332$ (water, IOS1 vs IOS2). |
| 4D | Two-way ANOVA: Interaction: $F_{(2,36)}=39.26$, $p<0.000$, PTM: $F_{(2,36)}=39.25$, $p<0.0001$<br>NPA: $F_{(1,36)}=5.108$, $p=0.030$. Bonferroni's multiple comparison test: nitro: $p=0.0005$, phospho: $p<0.0001$, total: $p>0.99$. |

## Results

### NO induces nitration at Y816 in TRKB receptors *in vivo* and *in vitro*

We first investigated whether TRKB might be nitrated at tyrosine residues *in vitro* and *in vivo.* By co-IP of nitroY and TRKB, we found basal levels of TRKB nitration in both cultured neurons and brain tissue of naïve mice (Fig 1). Nitric oxide donor sodium nitroprusside (SNP) (Fig 1A, 1B) and nitric oxide synthase substrate L-arginine (Fig 1C) increased TRKB nitration in cultured neurons. Since a previous computational analysis indicated Y816 residue as potential target of nitration (Biojone et al., 2015), we investigated if nitration is abolished in TRKB mutants carrying a Y816F substitution. In N2a neuroblastoma cell line transfected to express WT or mutated GFP-tagged TRKB, we found that nitration induced by L-arginine treatment (1.5 mM, 15 min) in the wild-type TRKB was completely abolished by Y816F mutation (Fig 1D). In order to ensure the specificity of the co-IP nitroTRKB assay, we designed Y816 nitrated and Y816 phosphorylated synthetic peptides, based on the C-terminal sequence of TRKB (Fig 1E), and compared the assay’s ability to detect those post-translational modifications. The anti-nitroY antibody specifically recognized the nitrated TRKB peptide (Fig 1F), nonspecific reaction from anti-nitroY antibody against non-nitrated TRKB (either phosphorylated or unmodified peptide) was almost undetectable.

### Nitration restricts TRKB phosphorylation and PLCγ signaling

Activation of TRKB by BDNF induces phosphorylation of its intracellular tyrosine residues, increases interaction with PLCγ, and increases complexity of dendritic branching (Atwal et al., 2000; Woo et al., 2019). Thus, we measured those parameters to elucidate if nitration functionally affects TRKB signaling. Pretreatment with SNP attenuated BDNF-induced TRKB phosphorylation (Fig 2A) in cultured cortical neurons, while NPA *per se* increased it (Fig 2B). Importantly, inhibition of guanylyl cyclase, an enzyme that mediates NO effects in many physiological conditions (Garthwaite, 2016), did not mimic NPA effect upon TRKB phosphorylation. Cortical neurons (7 DIV) were treated with ODQ 1μM (concentration sufficient to abolish GC-dependent cGMP production (Boulton et al., 1995; Garthwaite, 2016) for 30 min, and phosphoTRKB(Y816) was assessed by ELISA, but we did not find increased TRKB phosphorylation [mean±sem: 100±28 (vehicle) and 67.17±32 (ODQ), unpaired T test, t_9_=0.7475; p=0.47, data not shown]. Co-IP assay showed that SNP reduced BDNF-induced interaction between endogenously expressed TRKB and PLCγ in hippocampal neurons (Fig 2C). Interestingly, we also found reduced interaction between the endogenous PLCγ and the nitrated TRKB C-terminal synthetic peptide, when compared to the unmodified peptide (Fig 2D), which suggests that TRKB nitration could additionally impair PLCγ signaling independently of TRKB Y816 phosphorylation. Finally, in agreement with the biochemical data that suggests facilitated TRKB signaling by NO inhibition, the nNOS inhibitor NPA increased neuronal branching, evidenced by Sholl analysis (Fig 2 E, F, G). Pretreatment with the TRKB antagonist ANA12 0.1 μM [concentration sufficient to block TRKB phosphorylation (Casarotto et al., 2021)], prevented NPA-induced neuritogenesis.

### Nitration modifies TRKB conformation, facilitates its recognition by endocytic proteins and leads it to lysosomal degradation

The finding that nitration could further impair TRKB signaling independently of TRKB Y816 phosphorylation, as evidenced by the reduced interaction between PLCγ and TRKB C-terminal synthetic peptide, raised the possibility that nitration could significantly affect TRKB conformation. Thus, we used the PyMOL software to predict the effect of nitration at Y816 on solvent accessibility of TRKB C-terminal portion (796-820 aa). Compared to unmodified TRKB (Fig 3A, dashed line), nitration (blue line) substantially reduced the solvent accessibility close to Y816 residue (814-816), suggesting that it might bury those residues and hamper the interaction with partner proteins that use that particular segment as binding site. Additionally, nitration induced other smaller changes farther from Y816 (between residues 796-812) either increasing or decreasing solvent accessibility, compared to unmodified TRKB. Since Y816 is a well-known site of phosphorylation, we also evaluated the effect of this post-translational modification (PTM) on TRKB solvent accessibility (Fig 3A, red line). Intringuily, we noticed that although nitration and phosphorylation induce a strikingly different effect on the region close to Y816 (814-816), the phosphorylation-induced effects on the 796-812 region are indistinguishable from those induced by nitration. That observation led us to hypothesize that nitration and phosphorylation could induce either similar or distinctive effects upon interaction of TRKB with partner proteins, depending on the TRKB region required for each particular protein-protein interaction. With that in mind, we decided to test the effect of nitration on the interaction between TRKB and other partner proteins besides PLCγ.

It has been demonstrated that AP2M, an adaptor protein involved in clathrin-mediated TRKB endocytosis, recognizes TRKB and binds more avidly to its C-term region, at 796-820 sequence, upon phosphorylation (Fred et al., 2019). Thus, we checked how nitration would affect TRKB and AP2M interaction. We observed that SNP increased (Fig 3B) while NPA decreased (Fig 3C) the interaction of TRKB with AP2M in neuronal culture, suggesting that nitration increases TRKB:AP2M interaction. Both the nitrated and the phosphorylated TRKB C-terminal peptides were also found to interact more with the endogenous AP2M compared to the control peptide (Fig 3D). Finally, since TRKB ubiquitination has been shown to mediate its endocytosis and degradation upon phosphorylation (Murray et al., 2019), we tested if nitration would affect TRKB interaction with ubiquitin, and endocytosis. SNP increased TRKB ubiquitination (Fig 3E) and reduced TRKB positioning on the neuronal cell surface (Fig 3F). To understand the fate of nitrated TRKB after endocytosis, we measured the amount of TRKB in late endosomes and lysosomes. SNP increased the co-localization of TRKB with both the late endosomal marker Rab7 (Fig 3G) and with the lysosomal marker Lamp1 (Fig 3H), indicating that nitration directs TRKB to lysosomal degradation.

### nNOS inhibitor NPA facilitates plasticity in the visual cortex

Since our data suggests that nNOS inhibition facilitates TRKB signaling, and TRKB activation has been shown to be crucial in restoring juvenile-like plasticity in the visual cortex (Maya Vetencourt et al., 2008; Casarotto et al., 2021), we hypothesized that nNOS inhibition would also be able to induce plasticity in that experimental model. Peroral treatment with NPA for 3 weeks, followed by 7 days of monocular deprivation combined with NPA treatment, induced a shift in ocular dominance towards the non-deprived eye (Fig 4C) which was not observed in the control (water treated) mice, indicating increased plasticity induced by NO inhibition. In an independent cohort of naive mice (Fig 4D), 3 weeks of NPA treatment reduced the basal levels of TRKB nitration in more than 80%, while significantly increasing phosphorylation. The expression of TRKB seems to be unaffected by NPA treatment since no difference was found in the total levels of TRKB.

## Discussion

The idea of a functional interaction between NO and TRKB signaling has definitely been around for a while (Biojone et al., 2015; Stanquini et al., 2018; Ribeiro et al., 2019). However, very little is known about how that could happen at molecular and cellular level. We found that TRKB is a target of tyrosine nitration not only in neuronal cultures treated with NOS substrate L-arginine or NO donor SNP but, intriguingly, also in the brain tissue of healthy naïve mice. This evidence challenges the notion that nitration is a sign of physiological imbalance observed under oxidative stress; rather it shows that nitration of TRKB by nNOS takes place under physiological conditions, suggesting that nitration might play a significant role in modulating TRKB signaling in the healthy brain.

We found that NO directly targets Y816 residue in TRKB. Importantly, that residue is also a crucial starting point for many of TRKB-mediated effects since its phosphorylation, in response to BDNF, recruits the PLCγ pathway and regulates LTP (Minichiello et al., 2002), transcriptional regulation (Reichardt, 2006), dendritic arborization (Berghuis et al., 2006), etc. Although nitration and phosphorylation do not take place in the same carbon within tyrosine residue, those post-translational modifications are believed to be mutually excluding chemical reactions in a given tyrosine residue (Abello et al., 2009). Of interest, nitration shifts the pKa of the phenolic hydroxyl group, thus preventing phosphorylation by tyrosine kinases which act upon neutral phenolic hydroxyl group but not upon negatively charged phenolate (Abello et al., 2009). In this context, nitration of Y816 would be of high biological relevance due to its potential to restrict TRKB-PLCγ cascade, which led us to further explore that possibility. Indeed, we observed that NO donor SNP not only decreased TRKB phosphorylation of Y816 at basal levels, but also consistently reduced phosphorylation in response to increasing concentrations of exogenous BDNF. That evidence suggests that nitration is an important physiological regulator of TRKB receptors, being able to negatively affect the efficiency of TRKB signaling even if its ligand BDNF is freely available.

In line with decreased phosphorylation of TRKB at Y816, induced by SNP, we also found decreased interaction between PLCγ1 and endogenous TRKB. Importantly, decreased interaction was also found between PLCγ1 and Y816 nitrated peptide corresponding to the C-terminal portion of TRKB, suggesting that nitration impairs PLCγ cascade not only by preventing Y816 phosphorylation but also by further decreasing PLCγ binding to the nitrated TRKB site.

Besides anchoring PLCγ, Y816 TRKB residue is important as a recognition motif for AP2M, an adaptor protein, part of AP2 complex, involved in clathrin-mediated endocytosis of TRKB (Fred et al., 2019). We found increased interaction between AP2M1 and endogenous TRKB from SNP-treated neurons, and also between AP2M1 and the nitrated peptide corresponding to the C-terminal portion of TRKB. Furthermore, interaction of TRKB with the late endosomal marker Rab7 and the lysosomal marker Lamp1 was increased. Taken together, these data suggest that nitration of TRKB at Y816 marks TRKB to be endocytosed and degraded in lysosomes.

Intriguingly, the data on interaction of C-term.TRKB:AP2M1 and C-term.TRKB:PLCγ suggests that nitration differentially affects TRKB interaction with its partner proteins, thus mimicking phosphorylation regarding recognition by some - but not all – TRKB partner proteins. In general, tyrosine nitration introduces additional negative charge to the protein, which modifies the local charge distribution thus changing the protein conformation. Additionally, nitration introduces a bulky attachment to the protein which might contribute, by steric hindrance, to changes in conformational and interaction with partner proteins (Radi, 2013). We investigated, *in silico,* potential conformational changes in TRKB induced by nitration at Y816 and found a decrease in solvent accessibility close to this tyrosine residue, and some additional changes (either increasing or decreasing solvent accessibility at different points of the C-terminal sequence) up to 20 amino acids further from the PTM site. Interestingly, simulation of the effect of Y816 phosphorylation shows that both PTMs induce strikingly similar effects on the exposition of most aminoacids residues within the segment analyzed, except for the nitration-mediated burying of residues between 812-816, suggesting that nitration and phosphorylation might induce some similar, but also some opposite effects regarding TRKB interaction with partner proteins, depending on the protein segment required for each particular interaction.

Phosphorylation of TRKB and its fate have been well characterized by many groups over the years: upon BDNF binding, phosphorylated TRKB recruits adaptor proteins, such as clathrin-associated AP2 complex and ubiquitin ligase, resulting in TRKB internalization into endosome (Kononenko et al., 2017; Murray et al., 2019). Importantly, internalized phosphorylated TRKB is signaling competent, composing vesicles known as signalosomes, and initiates signaling through PI3K/Akt, Ras/ERK, and PLCγ pathways as it moves along the neurites (Harrington and Ginty, 2013). Similarly, tyrosine nitration in TRKB also recruits AP2 complex and triggers ubiquitination, thus inducing TRKB internalization and subsequent degradation through proteasome and lysosome pathways. However, differently from what happens upon phosphorylation, we showed that nitration repeals PLCγ binding, thus it does not initiate signaling through downstream pathways and further impairs the signaling by decreasing the availability of TRKB, on the neuronal surface, to its ligand BDNF. Altogether, our results suggest that the endogenous BDNF-TRKB signaling is facilitated when TRKB nitration is prevented.

Effective TRKB signaling, combined with proper environmental stimuli, has been pointed as crucial for the clinical recovery from many neurological and psychiatric illnesses by allowing the rewiring of disadvantageous connections (Umemori et al., 2018). We showed *in vitro* that inhibition of nitric oxide promotes neuronal branching, an effect that was prevented by the TRKB antagonist ANA-12. Moreover, using the ocular dominance shift in the visual cortex as a model, we found that plasticity can be induced by inhibiting nNOS-derived NO production, an effect associated with reduced TRKB nitration and facilitated TRKB activation in the same brain area. Nitric oxide production is regulated by neuronal activity, through calcium-calmodulin modulation of nNOS activation (Hardingham et al., 2013), thus placing NO as an ideal candidate to mediate environmental stimuli-induced changes in the circuitry. Moreover, nitrergic neurons are distributed throughout layers II-VI (Yousef et al., 2004) of the visual cortex, and some nitrergic neurons (type I cells) present long-range neurites, which might allow strategic local NO production to modulate the circuitry. Of importance for the present study, BDNF-TRKB signaling has been shown to modulate both natural critical periods of plasticity (Hanover et al., 1999; Huang et al., 1999) and experimentally induced plasticity (Lesnikova et al., 2020; Casarotto et al., 2021) in the visual cortex.

Activation of TRKB, and the pY816 in particular, has been shown to be critical in the mechanism of antidepressant drugs (Castrén and Monteggia, 2021). We recently showed that antidepressants act by directly binding to TRKB transmembrane domain dimers and thereby allosterically promoting BDNF signaling through phosphorylated Y816, leading to, among others, increased spine formation and reactivation of critical period plasticity in the visual cortex (Casarotto et al., 2021). In animal models, inhibition of nNOS has been shown to display antidepressant-like effects (Harkin et al., 1999; Stanquini et al., 2018; Joca et al., 2019). Our present results suggest that NO directly nitrates Y816 and thereby impedes phosphorylation of Y816, which is expected to inhibit antidepressant activity. Therefore, our data further emphasizes the role of TRKB as a critical nexus for antidepressant drugs action.

In conclusion, we propose that TRKB nitration could serve as a useful cellular tool to limit or select TRKB activation accordingly in many physiological neuronal processes. Moreover, since deregulation of nitric oxide production has been implicated in the etiology of many brain disorders (Tripathi et al., 2020) - including some in which TRKB signaling is impaired, such as stress-related disorders (Joca et al., 2019) - it would be of interest to further investigate the potential involvement of TRKB nitration in other pathophysiological contexts.

## Acknowledgements

The authors would like to thank Seija Lågas and Sulo Kolehmainen for their technical assistance. Funding for this study was provided by European Research Council (#322742 – iPLASTICITY), EU Joint Programme - Neurodegenerative Disease Research (JPND) project CircProt (#301225), Academy of Finland (#294710 and #307416), Sigrid Juselius Foundation, Jane and Aatos Erkko Foundation.

## Conflict of interest

The authors declare no conflicts of interest.

## Author contributions

CB and EC conceptualized the study; CB and PC designed and analyzed the experiments; CB, PC, CC, MF, RH, AL, MV collected data; CB wrote the first draft of the manuscript, all authors edited and approved the final version of the manuscript.

## Notes

### Competing Interest Statement

The authors have declared no competing interest.

